# Lysophosphatidylserine Nanoparticles Induce Oral Tolerance Independent of TIM-4 and Canonical PS Processing

**DOI:** 10.64898/2026.09.24.754158

**Authors:** Edwin J. Ovalle, Vincent Chak, Elizabeth A. Wohlfert, Sathy Balu-Iyer, Jason G. Kay

## Abstract

Protein replacement therapy is highly effective for treating many protein-deficiency disorders, but its long-term clinical benefit is often limited by anti-drug antibody responses that neutralize therapeutic proteins, compromising safety and efficacy. To address these unwanted immune responses, a lipid nanoparticle–based immunotherapy platform has been developed to induce antigen-specific tolerance. Mice treated with antigen associated with either double-chain phosphatidylserine (PS) or lysophosphatidylserine (LysoPS), a single-acyl-chain phospholipid, showed reduced antibody titers following subcutaneous pre-exposure. However, only LysoPS nanoparticles induced tolerance after oral administration, suggesting structural differences between PS and LysoPS can impact tolerogenic potential, depending on the site of delivery. As an apoptotic “eat-me” signal, PS is recognized by phagocytic receptors such as TIM-4 and is associated with LC3-associated phagocytosis. These nanoparticles have been hypothesized to promote “apoptotic mimicry” to regulate immunity. However, the contribution of TIM-4 to oral tolerance remains unclear. Here, we investigated the role of TIM-4 in LysoPS-mediated oral tolerance using a knockout mouse model. Our data showed that TIM-4^-/-^ mice induced tolerance to ovalbumin encapsulated in LysoPS comparable to that of wild-type mice. We also found that LysoPS nanoparticles did not follow the same surface engagement and intracellular trafficking pathways as PS nanoparticles. These results reveal that, despite sharing the same headgroup, differences in acyl chain number critically shape nanoparticle–immune cell interactions and provide a mechanistic framework that may explain the unique ability of LysoPS nanoparticles to induce oral tolerance.

## Introduction

Protein replacement therapy (PRT) has played an important role in treating many inherited diseases by providing recombinant therapeutic proteins to replace deficient or nonfunctional proteins [1–3]. However, repeated exposure to these recombinant therapeutic proteins can trigger unwanted immune responses, such as the development of anti-drug antibodies (ADA), which can reduce treatment effectiveness, as seen in hemophilia patients treated with factor VIII and Pompe disease patients treated with recombinant alpha-glucosidase (GAA) [4–7]. Similar unwanted immune responses to proteins, whether exogenous, endogenous, or dietary, also occur in allergies, type I diabetes, and celiac disease, respectively, highlighting the need for strategies that can induce antigen-specific tolerance [8–10]. As a potential strategy to address these unwanted immune responses, we developed a nanoparticle-based immunotherapy that encapsulated the target antigen of interest with either phosphatidylserine (PS) or lysophosphatidylserine (LysoPS) to induce antigen-specific tolerance [11]. Although both PS and LysoPS nanoparticles can induce tolerance after subcutaneous administration, only LysoPS nanoparticles successfully induce tolerance after oral administration, suggesting that the structural differences between PS and LysoPS are critical for promoting oral tolerance [12–14].

PS is a negatively charged phospholipid found in eukaryotic membranes and acts as an apoptotic marker [15, 16]. Under homeostatic conditions, PS is primarily located in the inner leaflet of the plasma membrane [17, 18]. When a cell undergoes apoptosis, PS is flipped to the outer membrane, increasing its surface exposure and allowing it to be recognized by antigen-presenting cells (APCs) (macrophages and dendritic cells (DCs)) as an “eat-me” signal [19–21]. APCs recognize PS via multiple surface receptors, including TIM-1, TIM-4, BAI1, stabilin-2, TAM, and members of the CD300 family [22–27]. We previously conducted a study that briefly examined the role of one PS receptor in mediating tolerance; the receptor TIM-4 was blocked with a functional antibody in vivo before antigen-nanoparticle exposure and a failure to induce LysoPS and PS nanoparticle tolerance was observed [11]. In addition, other groups using macrophages lacking the TIM-4 receptor have reported drastic reductions in the uptake of PS-containing bodies, despite the presence of other PS receptors, underscoring the importance of TIM-4 in mediating PS uptake [28, 29]. TIM-4 is a specialized PS receptor that mainly acts as a tethering molecule due to the lack of a cytoplasmic tail [29, 30]. TIM-4 surface binding is sensitive to the density of serine exposure on the bilayer surface, as increased surface exposure of PS on the outer leaflet of the plasmalemma of an apoptotic cell leads to stronger TIM-4 interaction [31]. In contrast, LysoPS, a derivative of PS containing a single acyl chain, is relatively understudied. LysoPS can exert multiple effects across different cell types, including histamine release [32], calcium flux [33], fibroblast migration [34], and nerve cell proliferation [35]; however, few studies have examined its mechanism of action. LysoPS’s single acyl chain structure affects nanoparticle formation. LysoPS nanoparticles are cone-shaped structures that prevent tight lipid packing in the nanoparticles, producing smaller, more tightly curved particles with greater LysoPS localization to the outer leaflet and increased surface serine exposure compared with larger, more rigid PS nanoparticles [36, 37]. We therefore hypothesized that LysoPS nanoparticles would interact stronger with TIM-4 than PS nanoparticles.

Internalization of PS-displaying bodies via PS receptors leads to a specialized uptake mechanism called efferocytosis, which enables rapid breakdown and recycling of cellular components [38–40]. During efferocytosis, phagosomes can use traditional autophagy-related proteins, including microtubule-associated protein light chain 3 (LC3), to form an LC3-associated phagosome (LAPosome) for degradation of apoptotic cells. Post-engulfment of apoptotic cells, APCs secrete anti-inflammatory cytokines interleukin-10 (IL-10) and transforming growth factor (TGF)-β, while simultaneously inhibiting the production of inflammatory cytokines tumor necrosis factor (TNF)-α, IL-1β, and IL-12 [41–43]. This process of dampening the immune response to apoptotic cells is important for preventing autoimmunity, as defects in apoptotic cell recognition have been observed in many autoimmune diseases [44, 45].

While TIM-4 plays an important role in PS-mediated immune tolerance and studies have shown that PS directly binds to TIM-4, the role of TIM-4 in LysoPS has not been investigated. In this study, we investigated the tolerogenic potential of LysoPS nanoparticles containing ovalbumin (OVA) to induce oral tolerance using TIM-4 knockout mice. We hypothesized that increased serine exposure on LysoPS nanoparticles would promote enhanced uptake and processing by APCs via TIM-4 and the LC3-associated phagosome pathway. However, to our surprise, TIM-4 knockout mice induced tolerance to OVA when encapsulated in LysoPS comparable to wild-type mice. These results suggest that although LysoPS and PS have identical headgroups, differences in the number of acyl tails lead to distinct uptake pathways that may explain differences in tolerance induction ability between the lipids observed during oral tolerance induction.

## Materials and Methods

### Materials

The following lipids were used: 1-oleoyl-2-hydroxy-sn-glycero-3-phospho-L-serine (sodium salt; 18:1 LysoPS, Cat. 326589-90-6), L-α-phosphatidylserine (BrainPS, Cat. 383907-32-2), and 1,2-dimyristoyl-sn-glycero-3-phosphocholine (DMPC; Cat. 18194-24-6) were purchased from Avanti Polar Lipids. DiI dye (Cat. D3911) was purchased from Invitrogen.). Endograde ovalbumin (OVA) was obtained from BioVendor LLC (Asheville, NC, USA). Solvents and buffer salts were purchased from Fisher Scientific (Fairlawn, NJ, USA). Antibodies for flow cytometry were obtained from eBioscience, Inc. (San Diego, CA, USA), BioLegend (San Diego, CA, USA), and Cytek Biosciences (Fremont, CA, USA). ELISA kits were purchased from Chondrex (Woodinville, WA, USA). Macrophage culture medium was composed of RPMI 1640 (Gibco), 2 mM L-glutamine (Corning), 10% fetal bovine serum (FBS) (Corning), 100X Antibiotic-antimycotic (Gibco), 1 mM Sodium pyruvate.

### Mice

B6.Cg-Timd4tm1Kuch/J (TIM4^-/-^, Strain #:035869) and background strain C57BL/6J (B6, Strain #:000664), were obtained from Jackson Laboratory. (Bar Harbor, ME USA) TIM4^-/-^ mice were genotyped to confirm strain identity before study initiation. Both male and female C57BL/6J mice aged 8–12 weeks and TIM4^-/-^ mice aged 30–70 weeks were used. All animal procedures were performed in accordance with protocols approved by the Institutional Animal Care and Use Committee (IACUC) at the University at Buffalo, The State University of New York. Endotoxin levels were monitored using the Endosafe Endochrome-K endotoxin assay kit (Charles River Laboratories, Charleston, SC, USA), and only buffers and formulations with endotoxin levels <0.05 EU/mL were used in vivo.

### Nanoparticles preparation

Phosphatidylserine or lysophosphatidylserine nanoparticles were prepared at a 30:70 molar ratio of Brain PS or LysoPS with DMPC using the thin-film method as previously described [46]. For fluorescently label nanoparticles, 30:69.5:0.5 molar ratio of Brain PS or LysoPS with DMPC and DiI was used. The film was rehydrated in 5 mM citrate buffer at pH 4.0 and extruded eight times through a polycarbonate membrane with 100 nm pore size using a high-pressure extruder (Mico, Inc., Middleton, WI). The final lipid concentration was confirmed via a standard phosphate assay [47]. Ovalbumin was loaded into the lipid nanoparticle at a lipid molar ratio of 1:1000 and incubated at 37°C for 30 minutes. The size of the nanoparticle was measured using a Nicomp 380 Submicron Particle Sizer. (Entegris, Santa Barbara, CA). Data were analyzed using the intensity-weighted Gaussian analysis.

### Oral tolerance study

C57BL/6J (n=8/group) and TIM4^-/-^ (n=5/group) mice were prophylactically treated with buffer, 1 μg of OVA protein, or 1 μg of OVA protein in the presence of LysoPS nanoparticles, once weekly for 8 weeks. Starting at week 5, mice were rechallenged weekly with 2 μg of free OVA subcutaneously, 24 hours after oral administration, for four weeks. Mice underwent a two-week washout period after the final rechallenge. All mice were sacrificed, and plasma was collected via cardiac puncture containing 10% v/v acid citrate dextrose (ACD). A schematic of the study design is presented in **Figure 5A**. The spleen was collected for OVA-specific IgG titer measurement and B-cell immunophenotyping.

### Anti-OVA antibody ELISA

All ELISAs performed in this study were purchased from Chondrex (Woodinville, WA) and followed manufacturer protocols.

### In vivo T and B Cell Characterization

A single cell suspension was harvested from tissues and stained with LIVE/DEAD Fixable Blue Dead Cell Stain Kit, for UV excitation (cat#L34962, Life Technologies Corporation), CD138 BUV395 281-2 50ug (cat #740240), Foxp3 R718 3G3 100ug (cat#567465),APC/Fire™ 810 anti-mouse CD4 Antibody (cat#100480), Brilliant Violet 785™ anti-mouse CD197 (CCR7) Antibody (cat#120127), Spark YG™ 570 anti-mouse/human CD45R/B220 Antibody (cat#103286), cFluor® B515 anti-Mouse CD8a – 100T (cat#R7-20545), cFluor® R659 anti-Mouse CD25 – 100T (cat#R7-20575), This flow cytometry result were obtained using Cytek Aurora 5 Laser UV/V/B/YG/R (64 + 3 Channel) (Fremont, CA) and data were analyzed using Cytek cloud.

### Intestinal Antigen Presenting Cells PS Receptor Panel

The small intestine was harvested from C57BL/6J and TIM4^-/-^ mice following the Lamina Propria Dissociation Kit (Miltenyi Biotec). Cells were resuspended into single-cell suspension, blocked with anti-mouse CD16/32 (Biolegend) then stained for the following anti-mouse antibodies, as required, in 1% BSA: F4/80 PerCP-Cy5.5 (Clone BM8 Cat:123128) (Biolegend), CD11c Pacific Blue (Clone N418 Cat:117321) (Biolegend), CD51 Biotin (Clone RMV-7 Cat:104103) (Biolegend), CD61 RB780 (Clone 2C9.G2 Cat:755845) (BD Biosciences), CD366 (TIM3) Pe/Cyanine7 (Clone B8.2C12 Cat:134009) (Biolegend), MerTK Pe/Dazzle-594 (Clone 2B10C42 Cat:151523) (Biolegend), TIM4 Alexa-647 (Clone RMT4-54 Cat:130007) (Biolegend). Secondary staining was done with Streptavidin Alexa Fluor-488 (Cat S11223) (ThermoFisher Scientific). Cells were then fixed and analyzed using Cytek Aurora 5 Laser UV/V/B/YG/R (64 + 3 Channel) (Fremont, CA). Data was analyzed using FlowJo (BD Biosciences), and a gating strategy was performed using the Fluorescence Minus One (FMO) method.

### Peritoneal Cell Isolation and Culturing

Peritoneal cells were isolated from WT or TIM4KO mice by peritoneal lavage with ice-cold 3% FBS in phosphate-buffered saline (PBS). If red blood cells were present, peritoneal cells were incubated with red blood cell lysis buffer. To primarily select macrophages, peritoneal cells were plated at 1×10^6^ cells/ml in a 24-well plate and incubated in culture medium for 2 hours at 37°C in a 5% CO_2_ environment. This allowed macrophages to adhere to the plate; nonadherent cells were removed by gentle washing with warm media. Cells were then used for downstream applications.

### Western Blot

Peritoneal macrophages were treated with OVA-NPs at different time points. At each time point, cells were washed once with cold PBS and lysed in RIPA buffer (50 mM Tris-HCl (pH 8.0), 150 mM NaCl, 1.0% NP-40, and 0.1% Sodium Dodecyl Sulfate) with Pierce Protease Inhibitor (ThermoFisher Scientific). Cell extracts were separated by 4–20% Mini-PROTEAN® TGX™ Precast Protein Gels (BioRad). After protein separation, the gel was transferred to a polyvinylidene difluoride (PVDF) membrane. The membrane was washed in TBST (20 mM Tris-base, 150 mM NaCl, and 0.1% Tween-20, pH 7.6) and blocked in 1% BSA/TBST. The membrane was cut at the 37 kD size marker (Precision Plus Protein, BioRad), and the bottom half of the membrane was incubated with rabbit anti-LC3A/B antibody (Cat. 4108S, CellSignal), and the top half was incubated with mouse anti-Actin antibody (Cat. 612656, BD Biosciences) overnight at 4°C. After washing in TBST the membranes were incubated with the secondary HRP-linked antibody for 2 hours (anti-rabbit and anti-mouse, respectively). Western blots were developed using an enhanced chemiluminescence substrate (Pierce ECL Western Blotting Substrate (ThermoFisher Scientific). Quantification of the bands from the immunoblots was performed using densitometry (FIJI).

## Results

### TIM-4 deficiency does not impair LysoPS-OVA–mediated oral tolerance

To investigate the role of the TIM-4 receptor in LysoPS-mediated oral tolerance, both C57BL/6 (n=8) and TIM-4^-/-^ (n=5) mice were continuously treated with buffer, free OVA, or LysoPS-OVA for 8 weeks. Beginning in week 5, all groups were subcutaneously rechallenged with free OVA, as illustrated in **Figure 1A**. C57BL/6 mice that received LysoPS-OVA showed a non-significant trend toward lower OVA-specific IgG antibody titers (p = 0.07) than mice that received free OVA **(Figure 1B)**. This trend is consistent with our previous study on OVA tolerance [12, 13]. Surprisingly, we observed no differences in OVA titers between TIM-4^-/-^ and C57BL/6 mice treated with LysoPS-OVA (p = 0.34), suggesting that TIM-4 may not be required for LysoPS-mediated oral tolerance. However, TIM-4^-/-^ mice treated with free OVA showed significantly lower (p = 0.03) OVA titers compared to C57BL/6 mice treated with free OVA, comparable to the titers observed with LysoPS-OVA treatment (p = 0.65). Together, these findings suggest that while TIM-4 may not be essential for the tolerogenic effect of LysoPS, TIM-4 deficiency itself may promote a more tolerogenic phenotype in response to free OVA.

**Figure 1.**
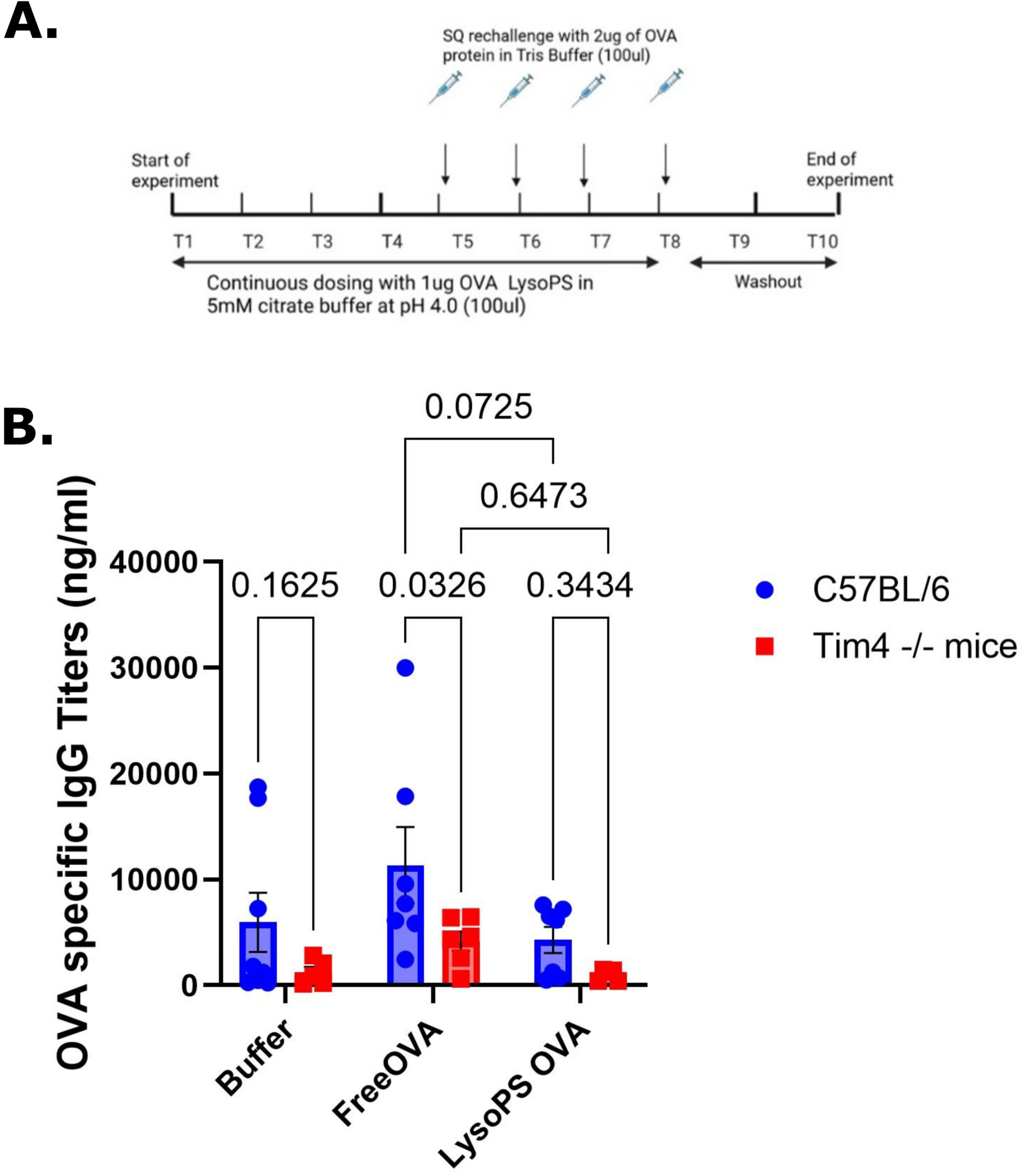
TIM-4 deficiency does not impair LysoPS-OVA–mediated oral tolerance. (A) Experimental design. C57BL/6 and TIM-4^-/-^ mice received continuous oral dosing of 1 µg OVA formulated with LysoPS in 5 mM citrate buffer (pH 4.0) from Week 1 to Week 6. At week 6, mice were subcutaneously rechallenged with 2 µg free OVA in Tris buffer. After a 2-week washout, blood was collected at the end of the study for antibody analysis. (B) OVA-specific IgG titers measured at the end of the study in C57BL/6 and TIM-4^-/-^ mice treated with Buffer, Free OVA, or LysoPS-OVA. Free OVA-treated C57BL/6 mice developed higher anti-OVA antibody titers compared with TIM-4^-/-^ mice, whereas LysoPS-OVA treatment reduced OVA-specific IgG responses in both groups. Data are presented as mean ± SEM. C57BL/6 mice had n=8/group while the TIM-4^-/-^ group had n=5/group. Statistical significance was determined using two-way ANOVA with Tukey’s multiple-comparisons test.

We further assessed the impact of LysoPS-OVA treatment on antibody-producing plasma cells. Spleens were collected at the end of the study from both mouse strains. Single-cell suspensions were prepared and stained for B220⁺CD138⁺ cells, which were quantified as plasma cells. C57BL/6 mice treated with LysoPS-OVA had a reduced frequency of splenic plasma cells compared with those treated with free OVA **(Figure 2)**. In TIM-4^-/-^ mice, free OVA and LysoPS-OVA treatments had lower plasma cell frequency than the buffer group. We also directly compared each treatment group between C57BL/6 mice and TIM-4^-/-^ mice. TIM-4^-/-^ mice treated with free OVA had significantly reduced splenic plasma cell frequency compared to C57BL/6 mice, similar to the frequency of LysoPS-OVA treatment. The significantly reduced IgG and plasma cell frequency in TIM-4^-/-^ mice compared with their wild-type counterparts, especially after free OVA administration, confirmed the tolerogenic phenotype in TIM-4^-/-^ mice.

**Figure 2.**
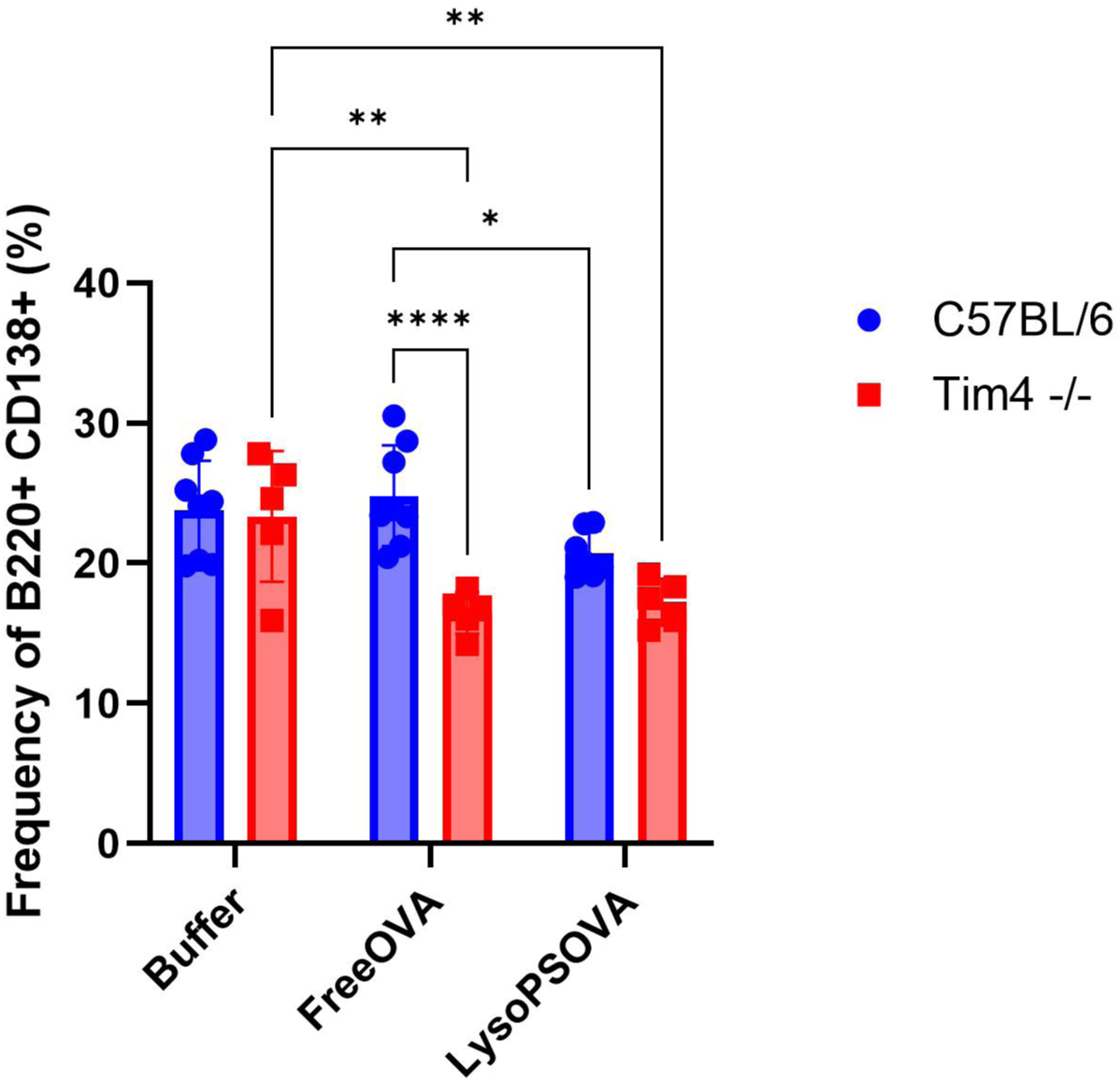
TIM-4^-/-^ mice show reduced plasma cell development. Frequency of B220⁺CD138⁺ plasma cells in spleens of C57BL/6 and TIM-4^-/-^ mice following treatment with Buffer, Free OVA, or LysoPS-OVA. Mice treated with Free OVA showed a significant reduction in plasma cell frequency in TIM-4-/- mice compared with C57BL/6 controls, whereas LysoPS-OVA treatment maintained similarly reduced plasma cell frequencies between groups. Data are presented as mean ± SEM. C57BL/6 mice had n=8/group while the TIM-4^-/-^ group had n=5/group. Statistical significance was determined using two-way ANOVA with Tukey’s multiple-comparisons test. *p < 0.05, **p < 0.01, ****p< 0.0001.

### TIM-4^-/-^ APCs do not compensate PS receptors for the lack of TIM-4

The immune system has evolved to have a redundancy of receptors for the clearance of apoptotic cells, with TIM-4 being one of 15 known PS receptors [48]. While TIM-4 is important for homeostasis and its knockout is not lethal, to date no study on TIM-4^-/-^ mice has determined whether these mice compensate for the loss of TIM-4 with other PS receptors. Given our in vivo results, with no difference in LysoPS treatment in TIM-4^-/-^ and wild-type mice, we speculated that TIM-4^-/-^ mice compensate for the absence of TIM-4 by upregulating other PS receptors. To test this, cells were harvested from the small intestine and gated for F4/80^+^ macrophages and CD11c^+^ DCs **(Figure 3A),** the expression of multiple PS receptors was assessed in both cell types **(Figure 3D)**. Intestinal TIM-4^-/-^ DCs showed no significant difference in expression of any tested PS receptors compared to wild-type, except for TIM-4 **(Figure 3E).** In TIM-4^-/-^ macrophages, there was a significant decrease in TIM-1 as well as TIM-4 **(Figure 3F).** In the absence of TIM-4, intestinal APCs do not compensate for the loss of PS receptors, a finding not previously reported. This provided additional evidence that LysoPS is not dependent on TIM-4 or other PS receptors.

**Figure 3.**
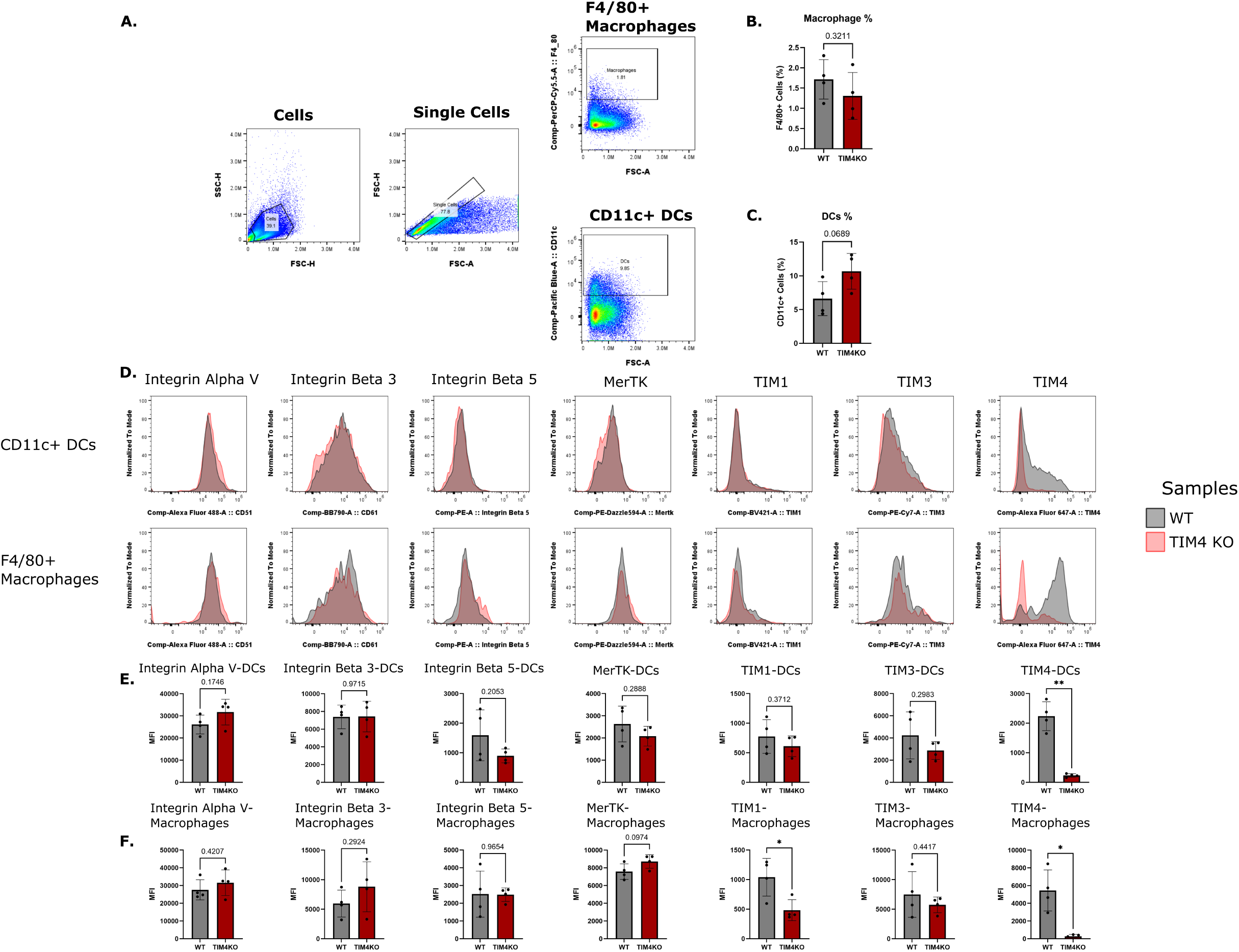
TIM-4^-/-^ APCs do not compensate PS receptors for the lack of TIM-4. Intestinal cells were isolated from wild-type or TIM-4^-/-^ mice. Cells were stained for multiple PS receptors after gating on macrophages (F4/80) and DCs (CD11c), as shown in the gating strategy (A). Frequency of (B) macrophages and (C) DCs in the small intestine. (D) Sample flow plots of PS receptors on intestinal macrophages or DCs are shown. The expression levels of the PS receptors were quantified in DCs (E) and macrophages (F). Data are presented as MFI expression, depicted as mean ± s.d. (n = 4) and analyzed using an unpaired t-test with Welch’s correction. *p ≤ 0.05; ns = not significant.

### LysoPS nanoparticles do not require the TIM-4 receptor for binding

The absence of TIM-4 has been shown to drastically decrease apoptotic cell uptake by macrophages [28, 29]. Because TIM-4^-/-^ APCs lack PS receptor compensation, we next assessed whether LysoPS nanoparticles required TIM-4 for binding/uptake. To test, we incubated lipid-encapsulated OVA nanoparticles with resident peritoneal macrophages from wild-type and TIM-4^-/-^ mice for 20 minutes to assess initial receptor-mediated binding and uptake. We included double-chain PS nanoparticles as a positive control, as they more closely resemble an apoptotic cell, and we speculated that nanoparticles made with both lipids would follow the same apoptotic mimicry pathway. There was a significant reduction in PS nanoparticle binding and uptake (as measured by DiI fluorescence) in TIM-4^-/-^ peritoneal macrophages compared with wild-type macrophages **(Figure 4A)**, as expected [28]. However, LysoPS nanoparticle fluorescence did not differ between TIM-4^-/-^ and wild-type macrophages **(Figure 4B)**. We also examined PS receptor expression in TIM-4^-/-^ peritoneal APCs to determine whether they had a similar phenotype to intestinal APCs, and we observed similar results **(Supplemental Figure 1).** The lack of TIM-4 on APCs affected the binding of PS nanoparticles, as previously reported, yet did not affect LysoPS nanoparticle binding, suggesting LysoPS nanoparticles may utilize a different mechanism from PS nanoparticles [28, 29].

**Figure 4.**
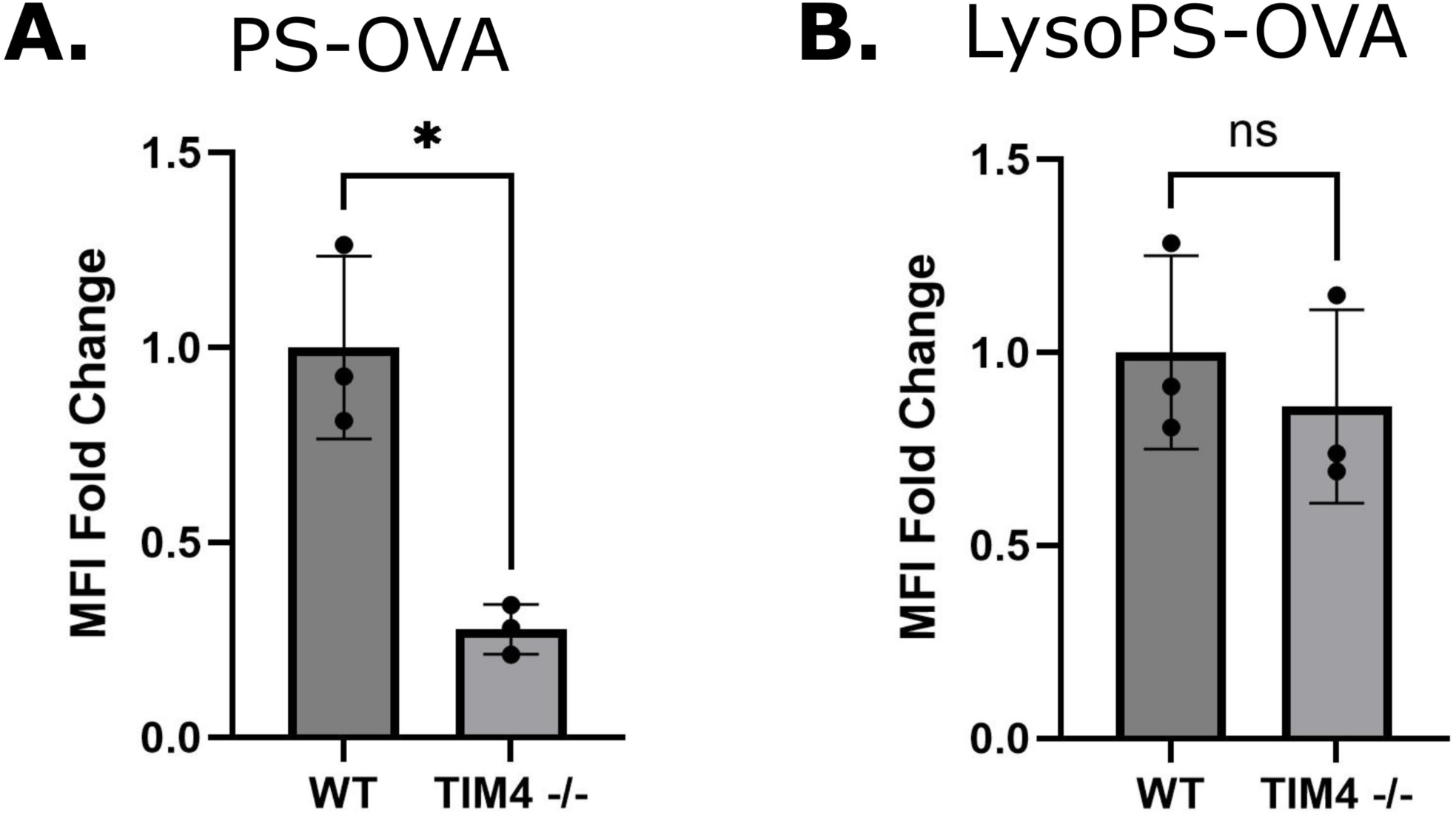
LysoPS nanoparticles do not require the TIM4 receptor for binding. Fluorescent OVA-NPs were incubated with peritoneal macrophages for 20 minutes. The expression of (A) PS-OVA and (B) LysoPS-OVA was recorded. Data is presented as MFI fold change, depicted as mean ± s.d. (n = 3). Data were analyzed using an unpaired t-test with Welch’s correction. *p ≤ 0.05; ns = not significant.

### LysoPS is not processed through LC3-associated phagocytosis

Although our data indicated that LysoPS nanoparticles were not internalized via the TIM-4 receptor, we aimed to investigate whether LysoPS nanoparticles are processed through the same pathway as PS. Apoptotic cells and PS-containing bodies are generally processed through the specialized LC3-associated phagocytosis (LAP) pathway, which is detectable by LC3 on the phagosome [43, 49]. During homeostasis, LC3, a regulator of autophagy, is present in the cytosol (LC3-I). When macrophages engulf apoptotic cells, LC3-I is lipidated (termed LC3-II) and binds to the phagosome. The conversion of LC3-I to LC3-II signifies autophagy activity, and the presence of LC3-II on a phagosome is indicative of LAP [50]. Peritoneal macrophages were treated with OVA-NPs for up to 20 minutes, due to LAP being short-lived on phagosomes [51]. Cell lysates were analyzed for LC3-I and LC3-II by western blotting **(Figure 5A)**. Macrophages incubated with PS nanoparticles showed an increase in the LC3-II/LC3-I ratio [52], peaking at 7.5 minutes, while macrophages incubated with LysoPS nanoparticles showed no changes in LC3 ratio **(Figure 5B, C)**. These results indicate that, in addition to LysoPS nanoparticles not requiring TIM-4 for uptake, they are not processed by LAP, and do not follow the same pathway as PS nanoparticles.

**Figure 5.**
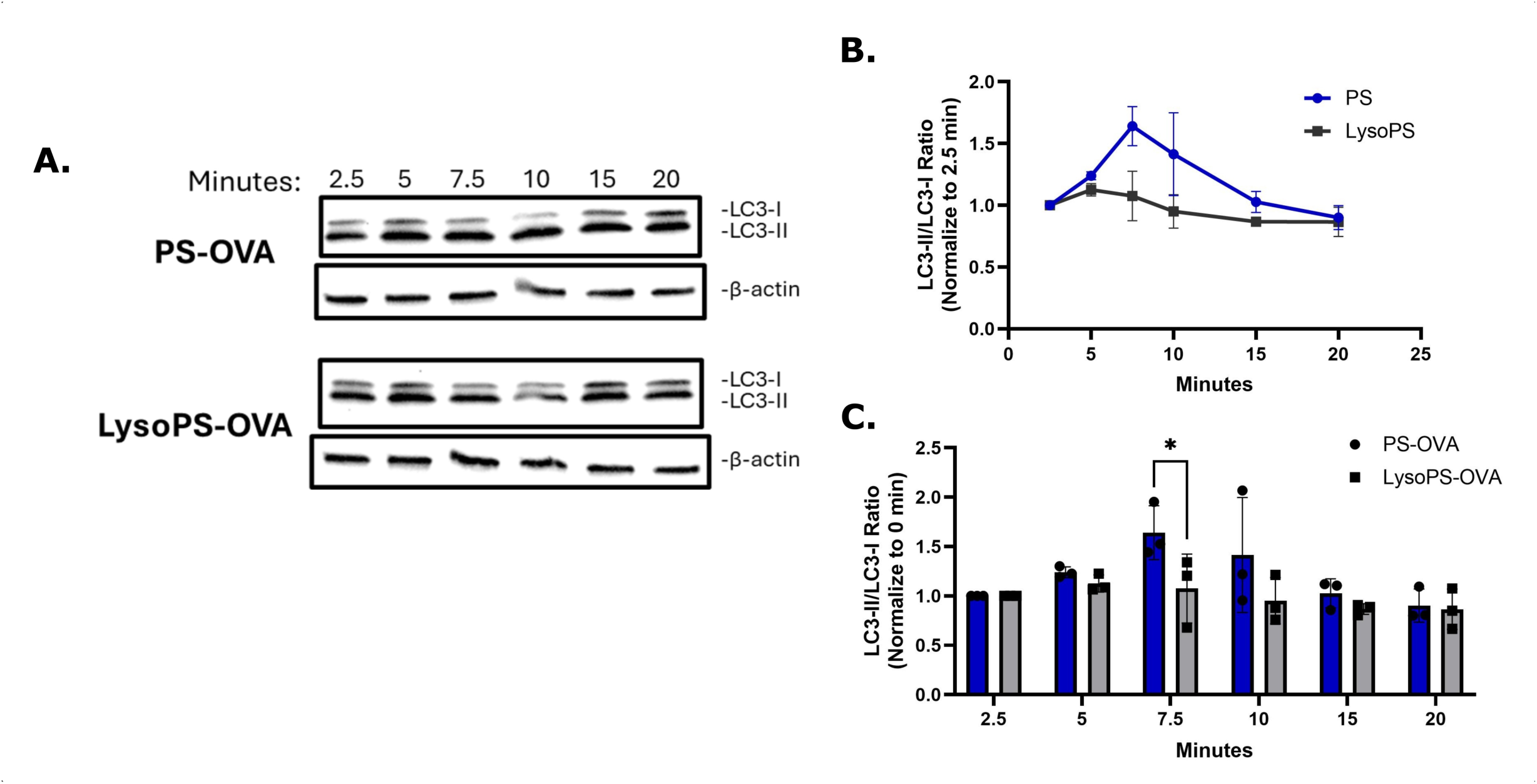
LysoPS-OVA is not processed through LAP. Sample western blots of LC3-I and LC3-II on PS-OVA or LysoPS-OVA treated peritoneal macrophages. The LC3-II/LC3-I ratio was calculated from band intensities, and the data were normalized to their respective 2.5-minute timepoints. Data are shown as mean ± s.d. (n = 3) in either a time-series plot (B) or a bar plot for the statistical test (C). Data were analyzed using an unpaired multiple t tests using Holm-Sidak’s multiple comparison test with a single pooled variance for all rows. *p ≤ 0.05; ns = not significant.

## Discussions

Previously, we have shown that both LysoPS and PS nanoparticles induce tolerance when administered subcutaneously [11, 46]. In addition, LysoPS, but not double-chain PS, induced antigen-specific tolerance when introduced orally in murine models [12], suggesting that the number of acyl chains on PS influences tolerogenic potential. The structure of PS impacts the biophysical characteristics of PS-containing nanoparticles, particularly the serine surface exposure, which is critical for immunological tolerance; incorporation of LysoPS into lipid nanoparticles increases the surface curvature of the vesicles, allowing more LysoPS to partition onto the nanoparticle surface (outer leaflet) than with double-chain PS [36].

TIM-4 is a PS receptor predominantly expressed on APCs, particularly macrophages and mature myeloid-derived DCs. TIM-4 contributes to immune regulation through multiple mechanisms: it acts as a ligand for TIM-1, promoting T-cell expansion and survival, and facilitates the engulfment of apoptotic cells via recognition of PS on the cell surface [28, 53]. TIM-4^-/-^ APCs showed drastic reductions in PS uptake, despite the presence of other PS receptors, highlighting the importance of TIM-4 for PS uptake [28, 29]. Furthermore, a previous study from our lab showed that subcutaneous pretreatment with a functional-blocking anti-TIM-4 antibody reversed both PS- and LysoPS-mediated tolerance, demonstrating that TIM-4 was essential for PS- and LysoPS-mediated immune regulation in peripheral tissues [11, 54].

In this study, we examined the importance of TIM-4 in LysoPS-mediated oral tolerance. In wild-type mice, LysoPS-mediated oral tolerance was consistent with our previous findings [12, 13], as shown by a slight, though not statistically significant, reduction in OVA-specific antibody responses. Meanwhile, LysoPS-OVA treatment had similar OVA titers in both TIM-4^⁻/⁻^ and wild-type mice, suggesting that LysoPS-mediated oral tolerance occurs independently of the TIM-4 receptor in gut-mediated tolerance. This was further confirmed in vitro, showing that LysoPS nanoparticles were taken up independently of TIM-4. In addition, we assessed the processing of LysoPS nanoparticles, initially expecting that both PS and LysoPS nanoparticles would follow the same mechanism, given that they share the same phosphatidylserine headgroup and differ only in the number of acyl chains. However, to our surprise, LysoPS nanoparticles were not processed like PS nanoparticles, suggesting that they follow distinct pathways.

These results contrast with our earlier studies that showed pretreatment with an anti-TIM-4 blocking antibody reversed PS-mediated tolerance induced by subcutaneous administration [11, 54]. Initially, we speculated that compensatory upregulation of other PS receptors may occur in the TIM-4^-/-^ model due to chronic loss of TIM-4, whereas in vivo TIM-4 antibody-blocking experiments acutely inhibit TIM-4 and may reduce the functional availability of PS receptors on APCs before compensation can occur. In contrast, TIM-4^-/-^ mice showed no overall compensation in other PS receptors on APCs for the receptors that we examined, a finding not previously reported. Despite the absence of TIM-4, the binding of LysoPS nanoparticles was not affected compared to double-chain PS nanoparticles.

Another group previously documented conflicting results between the TIM-4^-/-^ model and TIM-4 functional antibody-blocking experiments using a contact hypersensitization model [55]. A possible reason for the conflicting data in their case and our study may be that the functional TIM4-blocking antibodies have unknown off-target effects. Or, as we have shown here, an altered baseline phenotype in knockout immune cells and responses. A previous group also reported a shift in DC populations in TIM-4^-/-^ mice compared to wild-type mice [56]. In contrast to our data, the group that generated the TIM-4 knockout mice reported hyperactive T and B cells, elevated serum Ig levels, and the development of antibodies to double-stranded DNA due to TIM-4 deficiency [57]. Notably, the same study demonstrated that TIM-4 has a niche, compartment-specific role in apoptotic-cell clearance, as clearance was impaired in the peritoneal cavity but remained intact in the spleen. However, in our oral tolerance model, TIM-4^-/-^ mice exhibited a more tolerogenic phenotype, with reduced OVA-specific antibody responses and lower plasma cell frequencies than their wild-type controls. Further studies are needed to characterize TIM-4^-/-^ mice and determine how TIM-4 deletion affects them systemically. Alternatively, an inducible TIM-4 knockout model could minimize these systemic effects.

In addition to not requiring TIM4 for uptake, we also found that LysoPS nanoparticles are not processed through LAP. Post-engulfment of apoptotic cells can lead to this specialized processing, which promotes the production of anti-inflammatory cytokines, an important process for maintaining tolerance towards self and for inducing tolerance [41]. This form of non-canonical autophagy is characterized by LC3 conjugation to the phagosome [52]. Our results showed that PS nanoparticles induced a peak in LC3-II conversion at 7.5 minutes post-uptake. This processing rate is faster than in apoptotic cells [51], potentially because our nanoparticles differ in size from apoptotic cells. In contrast, LysoPS nanoparticles showed no LAP activity. LysoPS nanoparticles have been shown to induce tolerance here and in our previous studies [11, 12], suggesting that LAP is not required for tolerance induction by LysoPS. Traditional phagocytic/endocytic mechanisms could be an alternative pathway for macrophages to process LysoPS nanoparticles [58].

While LysoPS demonstrated the ability to induce antigen-specific tolerance, as with PS nanoparticles [11, 46] LysoPS and PS share similar structural components, except LysoPS contains only one acyl chain. We showed that this structural difference leads to LysoPS and PS following different uptake and processing mechanisms. In general, the role of LysoPS in vivo remains poorly understood, and though it is known to exert multiple effects across different cell types, including mast cell histamine release, neutrophil calcium flux, fibroblast migration, and nerve cell proliferation, little is known about its signaling mechanisms [59]. LysoPS has also been reported to have some effects in the context of apoptotic cell uptake, with apoptotic neutrophils and liposomes containing LysoPS having enhanced uptake by macrophages [60]. This enhancement was shown to occur via LysoPS interactions with a G protein-coupled receptor (G2A), activating the Rac1 pathway [61]. Currently, no other studies have examined LysoPS involvement in apoptotic clearance or inflammation. Our findings suggest that LysoPS may offer an advantage over PS in inducing oral tolerance due to its distinct mechanism of action.

### Overview, limitations, and future directions

LysoPS and PS nanoparticles elicit the same immunological response upon subcutaneous injection [11, 46], while following oral delivery, LysoPS, but not PS nanoparticles, is effective at promoting tolerance [12]. One receptor of interest involved in tolerance is TIM-4. In this study, we observed that LysoPS oral tolerance was not mediated through the TIM-4 receptor in a knockout model. However, in our in vivo model, we observed significantly reduced plasma cell production, we observed only a non-significant trend toward lower OVA titers with LysoPS treatment, consistent with our previous studies [12, 13]. OVA alone is a well-established antigen for inducing oral tolerance; however, in standard C57BL/6 mice, it is generally a weak oral immunogen unless paired with an inflammatory adjuvant or an OT-II adoptive transfer model that increases the detectable pool of OVA-specific CD4+ T cells [62–64]. This may explain why we did not reach significance in our OVA oral tolerance model. Another limitation is the substantial age difference between the C57BL/6J mice (8–12 weeks) and TIM-4^-/-^ mice (30–70 weeks), which may also have contributed to variability in the in vivo response. We found that LysoPS and PS nanoparticles utilize different binding/processing mechanisms within peritoneal macrophages, indicating a difference in mechanism of action between the nanoparticles and a potential explanation for how LysoPS outperforms PS during oral administration. Future studies should evaluate the role of the known LysoPS receptor, G2A, in tolerance induction and the traditional phagocytic mechanism. In addition, we did not examine LysoPS effects on DCs, the other major APC population. Macrophages were the focus here because they are the surveillance immune cells in the gut that efficiently sample and respond to their environment and are often the first APCs to interact with an antigen. Future studies will determine whether LysoPS and PS nanoparticles exhibit similar or distinct mechanistic effects when interacting with DCs.

## Conclusions

In conclusion, this study found that LysoPS nanoparticles do not follow the apoptotic mechanism as PS nanoparticles do, despite their shared serine headgroup [23]. LysoPS nanoparticles successfully induced tolerance in the absence of TIM-4, whose binding was not dependent on or processed by LAP. This study highlights differences between LysoPS- and PS-mediated interactions and provides insight into the mechanism of LysoPS oral tolerance.

## Acknowledgments

We acknowledge the UB Optical Imaging and Analysis Facility.

## Abbreviations

LysoPS: Lysophosphatidylserine
PS: Phosphatidylserine
APCs: antigen-presenting cells
DCs: dendritic cells
OVA: ovalbumin
LC3: light chain 3
LAP: LC3-associated phagocytosis

## Sources Funding

This work was supported by the National Institute of Allergy & Infectious Disease [grant number R01 AI169296] to SVB, JGK and EW. The content is solely the responsibility of the authors and does not necessarily represent the official views of the National Institutes of Health.

## Authorship Contribution Statement

EJO performed in vitro experiments, analyzed the data, prepared figures, and wrote the manuscript. VC performed the in vivo experiments, analyzed the data, and prepared figures. JGK, SVB and EW reviewed the manuscript and supervised the study.

## Disclosures

The authors declare no conflict of interest.

## Supporting Information

This manuscript contains supporting information.

---

**Supplemental Figure 1. TIM4KO Peritoneal Cells minimally alter the expression of other PS Receptors.**

Peritoneal cells were isolated from wild-type or TIM-4 -/- mice. (A) Cells were stained for multiple PS receptors after gating for macrophages (F4/80) and DCs (CD11c) following the gating strategy in. (B) Sample flow plots of PS receptors on peritoneal macrophages or DCs are shown. The expression levels of the PS receptors were quantified in (C) DCs and (C) macrophages. Data are presented as MFI expression, depicted as mean ± s.d. (n = 4-7) and analyzed using an unpaired t-test with Welch’s correction. *p ≤ 0.05; ns = not significant.

## Notes

### Competing Interest Statement

The authors have declared no competing interest.

