## Supplementary figures and images for "Lysophosphatidylserine Nanoparticles Induce Oral Tolerance Independent of TIM-4 and Canonical PS Processing"

### Supplemental Figure 1

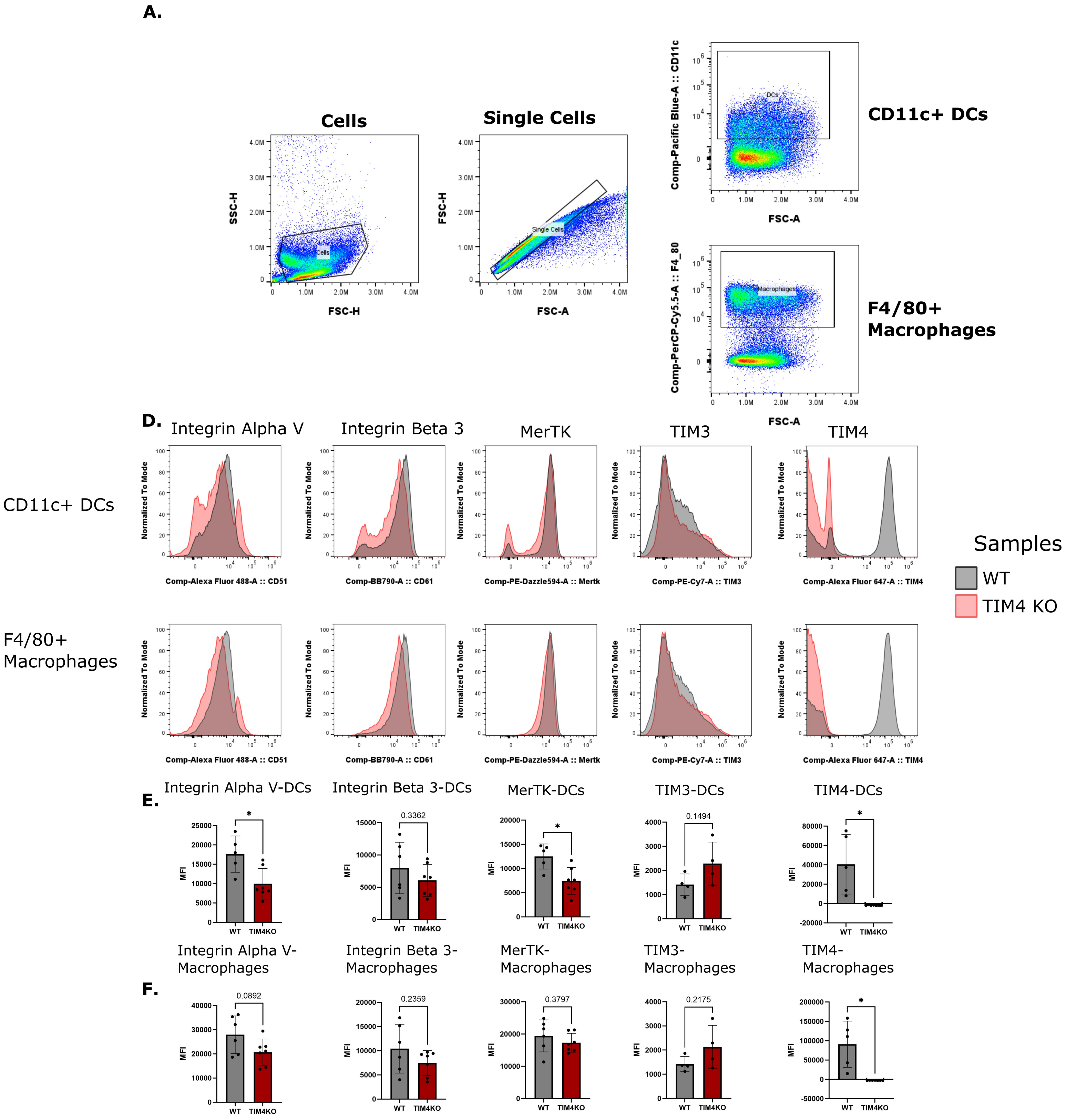
